# Generation of human chambered cardiac organoids from pluripotent stem cells for improved modelling of cardiovascular diseases

**DOI:** 10.1101/2021.05.21.445153

**Authors:** Beatrice Xuan Ho, Jeremy Kah Sheng Pang, Qian Hua Phua, Lee Chuen Liew, Boon Min Poh, Ying Chen, Yuin-Han Loh, Omer An, Henry He Yang, Veerabrahma Pratap Seshachalam, Judice LY Koh, Woon-Khiong Chan, Shi Yan Ng, Boon-Seng Soh

## Abstract

Recent progress on murine and human cardiac organoids have provided understanding to the developmental processes of the heart. However, there is still an unfulfilled need for improved modelling of cardiovascular diseases using human cardiac organoids. Herein, we report successful generation of intrinsically formed human chambered cardiac organoids (CCO) and highlight its utility in modelling disease. Single cell transcriptomic profiling of CCOs showed appropriate cardiovascular cell type composition exhibiting improved maturation. Functionally, CCOs recapitulated clinical cardiac hypertrophy by exhibiting thickened chamber walls, reduced ejection fractions, increased myofibrillar disarray and tachycardia. Therefore, CCOs improve current capabilities of disease modelling as an *in vitro* model bridging the gap to *in vivo* models, with the ability to assess functional parameters that previously can only be achieved in animal systems.

**One sentence summary:** Modelling cardiac hypertrophy using chambered cardiac organoids derived from human pluripotent stem cells.

## Main Text

Culturing of human pluripotent stem cells (hPSCs) derived tissues in 3D enables accurate *in vitro* research to recapitulate the microenvironment and geometric features of *in vivo* tissues (*1*). Novel innovations into 3D culture systems for various tissues make use of scaffolding, micro-engineering, and spheroidal aggregations (*2*). The self-organization capabilities of pluripotent cells are tapped into by organoid generation technology to promote embryonic-like development *in vitro* without the use of external scaffolds (*3*). These tissue specific organoids can be reproducibly generated, enabling high-throughput screening for developmental biology and translational research (*4-7*). In the context of cardiovascular biology, while progress has been mainly achieved for scaffold-based systems (*8, 9*), mouse cardiac organoids have been shown to be formed intrinsically, suggesting that a similar human system has the potential to be developed (*10, 11*). In the same vein, another research group recently published that human embryoid bodies coaxed towards to cardiovascular lineage can generate cardiac organoids resembling early embryonic cardiogenesis (*12*). However, there is still a need for an effective 3D cardiac organoid system that can appropriately represent majority of adult heart complications of structural cardiomyopathies and screen compounds at high-throughput. Current drug cardiotoxicity screens rely on the hERG assay, which is a useful determinant for adverse effects on cardiac electrical signaling, but fails to deliver any information on adverse mechanical effects on the heart (*13*). Henceforth, the current study details the generation of an effective Chambered Cardiac Organoid (CCO) model which fills up this gap in disease modelling. CCOs are shown to be composed of the appropriate cardiovascular cell types that exhibited improved maturation. Improved disease modelling was achieved by utilizing the functional chamber to model mechanical changes in ejection fraction and heart wall thickness, while also providing information on structural disarrays and arrhythmic tendencies at high-throughput.

Other organoid protocols published have demonstrated that progenitor-equivalents are necessary for the differentiation and self-organization in organoids (*4, 14-16*). As such, we reasoned that the proliferative ISL1^+^ cardiovascular progenitor (CVP) cells have the differentiation potential to generate the appropriate populations of cardiovascular lineages and enable the self-organization of CCOs. Tracking the lineage gene expression in hPSCs through 9 days of cardiomyocyte (CM) directed differentiation (*17*) showed the progression of the hPSCs leaving pluripotency (D0 to D4) and entering the mesodermal lineage (D1 to D6), with a concomitant upregulation of cardiovascular mesodermal and progenitor genes (D4, peaking by D7), before CM markers start to be expressed (D5 onwards) (Fig. 1, A and B). Therefore, the monolayer 2D CM differentiation protocol was adapted to include an organoid generation step and subsequent progenitor cell expansion at D7 (Fig. 1C). Matrigel was used to integrate CVPs and CMs within the CCOs at the early stage of formation. As Endothelin-1 (EDN1) has been shown to play an important role in expanding cardiovascular progenitor (CVP) cells *in vitro*, and transplanting early passages of the expanded CVP cells results in the formation of chamber-like structure *in vivo* (*18*), CCOs were cultured in EDN1 containing cardiosphere growing media for 7 days followed by maturation in spinner flask culture using RPMI/B27+ insulin media. The intrinsic formation of the chamber was tracked using brightfield imaging from D8 through D17, observed as a hollow core of a lighter shade present in the middle of the CCOs (Fig. 1D). A minimal cell number is required for chamber formation as organoids generated with 500k D7 cardiac differentiating cells was most robust in obtaining CCOs compared to 300k and 400k cells (Fig. 1E; movie S1). Cryosection and immunocytochemistry of D21 CCOs showed cardiomyocyte (CTNT) and smooth muscle (SMMHC) organization around the chamber (Fig. 1F). Low levels of Ki67+ cells indicated the expected low proliferation rates of matured CMs (Fig. 1F). Of note, the intrinsically formed hollow chamber was not a consequence of apoptotic cells as the low levels of cleaved Caspase-3 staining indicated healthy, non-apoptotic CMs (Fig. 1F). Furthermore, as chamber formation was observed by Day 8, it is unlikely a consequence of apoptotic cells in the organoid core (Fig. 1D).

**Fig. 1.**
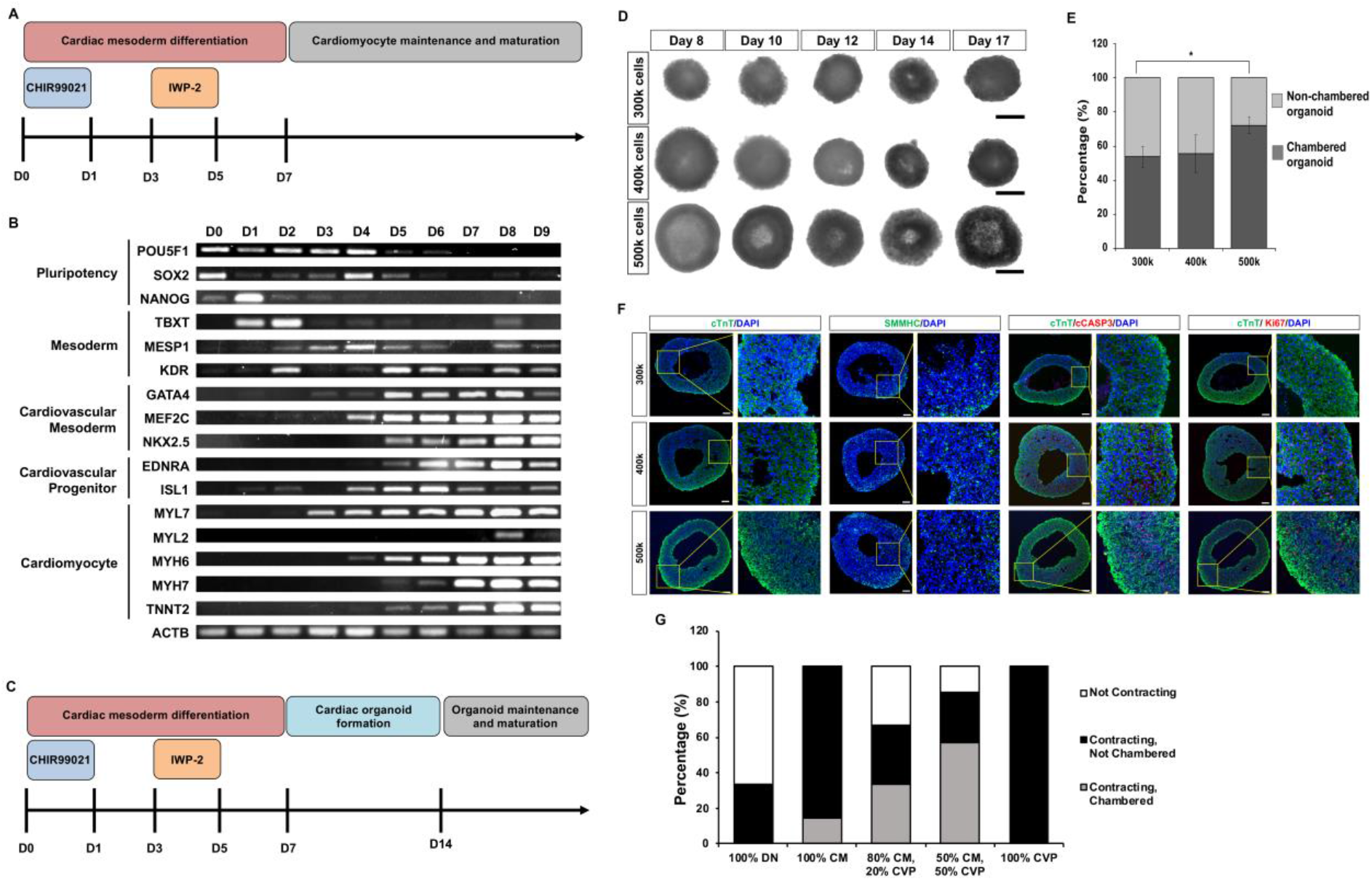
Generation and validation of human pluripotent stem cell-derived chambered cardiac organoids. (**A**) Schematic of the established hPSC-CM differentiation protocol. (**B**) RT-PCR tracking of relevant markers of cardiomyocyte differentiation from day 0 to day 9. (**C**) Schematic of the altered differentiation protocol to generate cardiac organoids. (**D**) Representative bright field images of CCOs generated using 300k, 400k or 500k cells from day 8 to day 17. (**E**) Percentage of CCOs generated using 300k, 400k or 500k cells per organoid (n = 8 organoids per group from 3 experiments; mean ± s.d; \**P* = 0.0141; two-tailed Student’s *t*-test). (**F**) Confocal immunofluorescence image of cardiomyocyte marker CTNT, smooth muscle cell marker SMMHC, proliferation marker Ki67 and apoptosis marker cleaved Caspase-3 in day 21 organoids generated from 300k, 400k or 500k cells. Scale bar, 100µm. (**G**) Percentages of contracting and chambered cardiac organoids generated using different compositions of cell populations. **DN**: Double Negative Cells; **CM**: SIRPA^+^ EDNRA^-^ Cardiomyocytes; **CVP**: SIRPA^+^, EDNRA^+^ Cardiovascular Progenitor Cells (n = 6 per group).

To further validate our hypothesis that CVPs are necessary for the intrinsic chamber formation, D7 CMs and CVPs were isolated using SIRPα (*19*) and EDNRA (*18*), respectively. SIRPα^-^/EDNRA^-^ double negative cells (DN), SIRPα^+^/EDNRA^-^ CMs and SIRPα^+^/EDNRA^+^ double positive CVPs were collected from the sort (fig. S1). Five compositions of organoids were generated [(1) 100% DN; (2) 100% CMs; (3) 80% CMs, 20% CVPs; (4) 50% CMs, 50% CVPs; (5) 100% CVPs], cultured to D21 and classified into three categories – not contracting, contracting without chamber formation, and contracting with chamber present (Fig. 1G). As expected, DN organoids of condition 1 either did not contract (67%) or contracted weakly (33%), while CM organoids of condition 2, were all contracting but most lacked chambers (86%). Only in the conditions where both CVPs and CMs were present did the cardiac organoids contracted and formed chambers, 33% and 57% of condition 3 and 4 respectively. Surprisingly, CVP organoids of condition 5 were all contracting with no chamber present, suggesting that not only are CVPs crucial in the formation of CCOs, there also exists an optimal balance between CVPs and CMs for sufficient cell-to-cell signaling for intrinsic chamber formation.

Transcriptome profiling using single-cell RNA sequencing showed that CCOs contained the appropriate cell composition and achieved improved maturation across a time course from D8 (right after CCO formation) to D14 (after CVP expansion) followed by D21 (after 1 week of maturation) (fig. S2; D8: 2567 cells, D14: 3585 cells, D21: 1946 cells). Highly variable genes were used to identify the top 20 principal components for unsupervised clustering and Uniform Manifold and Projection (UMAP) non-linear dimensionality reduction, and 10 major cell clusters were assigned identities based on gene markers expressed (Fig. 2, A and B, data S1) (*20*). CM clusters of each time point were identified using CM markers (*TNNT2*^Hi^/*ACTN2*^Hi^/*TTN*^Hi^). A cardiac mesodermal cluster were identified only in D8 and D14 CCOs with cardiogenesis related *SFRP2*, and other early mesodermal genes. An endothelial cluster containing cells from all 3 time points was recognized using endothelial markers (*CDH5*^Hi^/*SOX17*^Hi^/*KDR*^Hi^). An early cardiac fibroblast cluster and a late cardiac fibroblast cluster from D14 and D21 respectively was classified based on increasing expressions of cardiac fibroblast markers (*THY1*/*TCF21*/*SOX9*/*POSTN*), with some cells in the cluster also expressing epicardial markers (*WT1*/*SEMA3D*/*TBX18*). A cluster of mesendodermal cells were identified in D8 and D14 CCOs (*TBX3*^Hi^/*FOXA2*^Hi^), that likely progressed to a cluster of definitive endodermal cells present only in D21 CCOs (*AFP*^*+*^/*TTR*^*+*^/*VIL1*^*+*^/*FOXA2*^*+*^). A cluster of dividing G2/M phase cells was only found in D8 and D14 CCOs, highly expressing cell cycling genes (*MKI67*^Hi^, *CDK1*^Hi^). The cardiovascular lineages all exhibited increase in lineage specific gene expression from earlier to later time points (fig. S3). Immunofluorescence staining of dissociated organoids confirms the presence of the key cardiovascular cell types – CMs (CTNT), smooth muscle cells (SMMHC) and endothelial cells (CD31) (Fig. 2C). Quantitative gene expression comparisons to CMs maintained in 2D showed that smooth muscle (*SMA*) and vascular (*CD31, VCAM-1*) genes were upregulated in CCOs by D14 (Fig. 2D). By D21, a more pronounced increase in expression levels of CM (*TNNT2*), smooth muscle and vascular genes was observed (Fig. 2E). These findings indicate that not only do CCOs maintain the cardiovascular lineage cell types, but these supporting cell types also continue to stay viable and contribute to the maturation in the 3D CCO microenvironment.

**Fig. 2.**
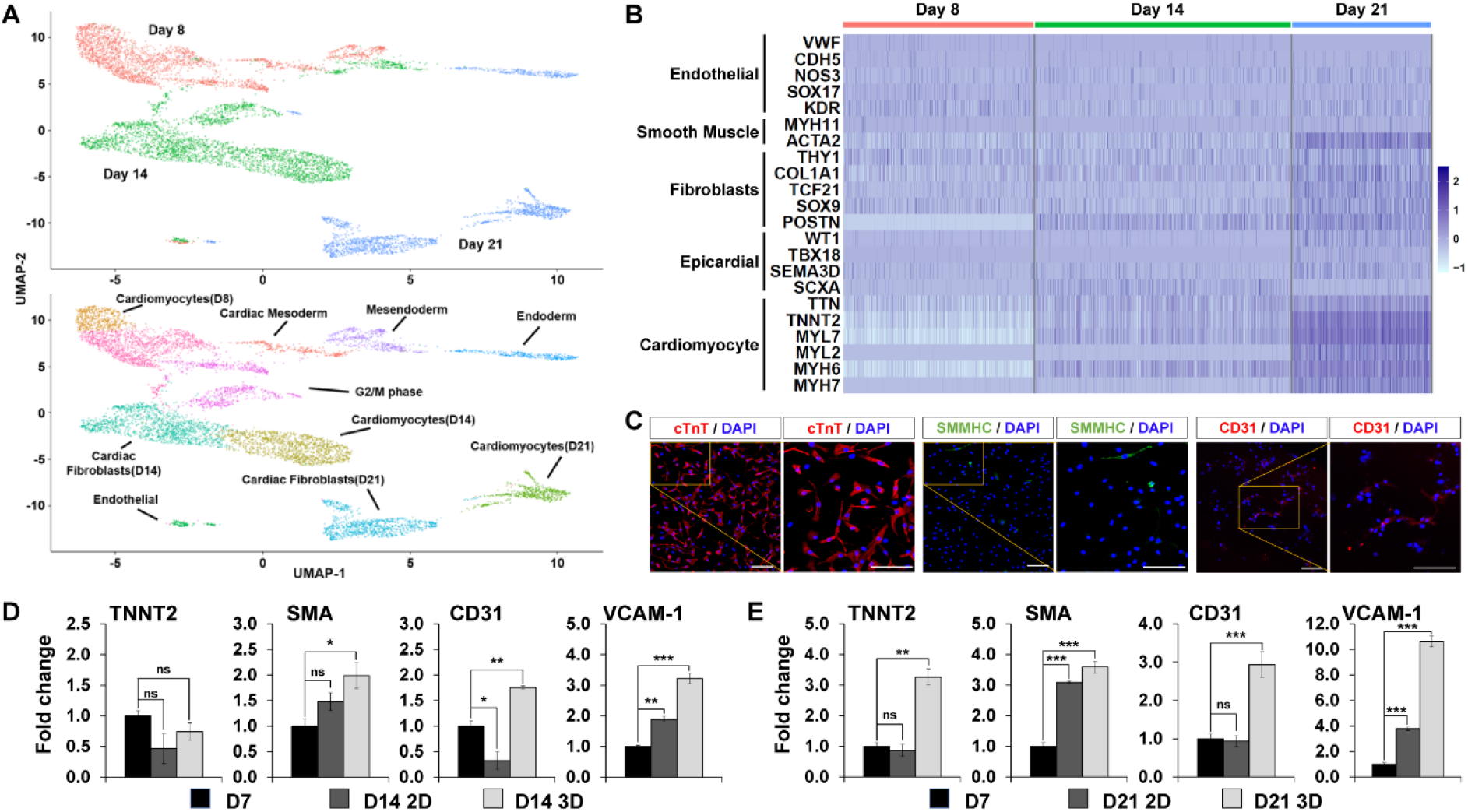
Chambered cardiac organoids is composed of cell types from the various cardiac lineages. (**A**) Uniform manifold approximation and projection (UMAP) of all cells, representing distinct identities of cells in day 8, day 14 and day 21 CCOs (top). UMAP revealed ten main cell clusters within the three time points (bottom). (**B**) Heatmap showing cell type specific genes expression differentially across time points. (**C**) Representative confocal immunofluorescence image showing expression of cardiomyocyte, smooth muscle cell and endothelial cell lineage specific markers in the dissociated CCOs. Scale bars, 100µm. (**D** and **E**) Quantitative gene expression comparison of 2D cultured cardiomyocytes and 3D cardiac organoids of day 14 and day 21 respectively, normalized to day 7 cardiomyocytes. Genes shown represent markers of cardiomyocyte, smooth muscle and endothelial cell lineages (n = 3; mean ± s.e.m; housekeeping gene, *ACTB;* \*\*\**P* < 0.001; \*\**P* < 0.01; \**P* < 0.05; two-tailed Student’s *t*-test).

Regarding the maturation of resident CMs, markers associated with different aspects of CM maturation, corresponding to progenitors, glycolysis, β-oxidation, CM specificity, and ion channels in the CM clusters over time were analyzed (Fig. 3A). As expected, there is maintenance of CVP genes *MEF2C* and *ISL1*, and an increase in *NKX2.5* from CVP enrichment between D8 through D14 (Fig. 3B). After switching out to maturation media, progenitor gene expressions decreased while *MEF2C* and *NKX2*.*5*, known to maintain CM identity, continued to be expressed (*21*). *EDNRA* expression did not change over the time points, suggesting that while it marks the initial pool of CVPs together with ISL1 expression, EDNRA expression is still necessary for continued interactions with EDN1 and vascular organization. CM maturation is linked with a metabolic switch from glycolysis to β-oxidation, and therefore glycolysis regulators *PGAM1* and *PGK1* were downregulated, while an upregulation of β-oxidation regulators *HADHA, HADHB* and *ACADVL* was observed across the time points (Fig. 3C). Across the board, CM specificity markers increase across the time points, with late maturation markers *MYL2* and *MYH7* expressed only at D21 (Fig. 3D). A similar pattern was observed for ion channel genes with some markers only expressed in the later time points (Fig. 3E). Quantitative gene expression comparisons to CM maintained in 2D showed expected results. By D14, while CCOs expressed increased β-oxidation markers *ACADVL* and *EHHADH*, CM specificity and ion channel markers are more significantly upregulated in 2D CMs, as the CCOs are being enriched for CVPs (Fig. 3F). By D21, all β-oxidation, CM and ion channel markers are highly upregulated in CCOs, highlighting marked improvements to maturation status compared to equivalent age 2D CMs (Fig. 3G).

**Fig. 3.**
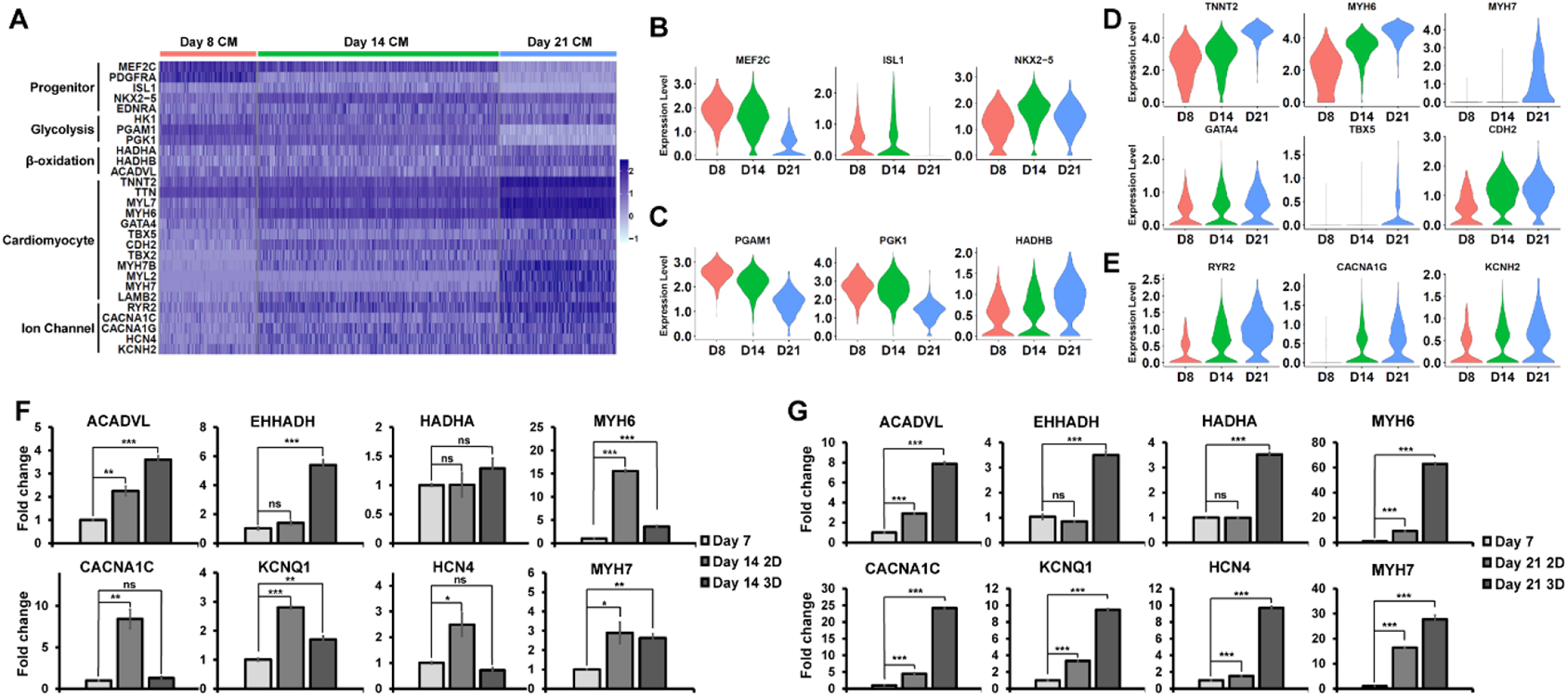
Chambered cardiac organoids improve maturity of resident cardiomyocytes. (**A**) Heatmap depicting the increased maturation of cardiomyocyte clusters within the CCOs over the time points. (**B-E**) Violin plots highlighting the specific genes corresponding to the changes in maturity of the cardiomyocyte clusters from day 8 (Red) to day 14 (Green) to day 21 (Blue). (**B**) Progenitor *MEF2C, ISL1, NKX2*.*5* expression. (**C**) Glycolytic *PGAM1, PGK1* and β-oxidation *HADHB* expression. (**D**) Cardiac *TNNT2, MYH6, MYH7, GATA4, TBX5* and *CDH2* expression. (**E**) Ion channel *RYR2, CACNA1G* and *KCHN2* expression. (**F** and **G**) Quantitative gene expression comparison of 2D cultured cardiomyocytes and 3D cardiac organoids of day 14 and day 21 respectively, normalized to day 7 cardiomyocytes. Genes shown represent the pathways of β-oxidation, ion channel and cardiomyocyte markers (n = 3; mean ± s.e.m; housekeeping gene, *ACTB;* \*\*\**P* < 0.001; \*\**P* < 0.01; \**P* < 0.05; two-tailed Student’s *t*-test).

Finally, to demonstrate functional utility of the CCO model, EDN1 was used as a cardiac hypertrophy (CH) inducer. Dysregulation of EDN1, a known vasoconstrictor has been shown to cause CH and other cardiomyopathies (*22-25*). To model CH, D28 CCOs were treated with two concentrations of EDN1 (50 and 100ng/ml) and their contraction parameters assessed over 3 weeks of treatment (Fig. 4A). CCO contraction videos were processed to track the boundary of the entire organoid (outer area) and inner chamber (inner area) to compute chamber wall thickness (Fig. 4B, fig S4, movie S1-3). While there were minor increases in the chamber wall thickness in control and 50ng/ml EDN1 treatment, only in the 100ng/ml EDN1 treatment was it sustained over three weeks (∼30-40%) (Fig. 4C). Analysis of the ejection fraction of the CCOs on the third week of treatment confirmed a significant ∼30% decrease in the 100ng/ml treatment group (Fig. 4D). The organoid contraction rate and rhythm were analyzed to be used as a measure of arrhythmic tendencies. Contraction frequency increased significantly in the 100ng/ml treatment group, doubling by week 3, while contraction variability decreased, correlating with the expected tachycardiac phenotype observed in clinical settings (Fig. 4, E and F).

**Fig. 4.**
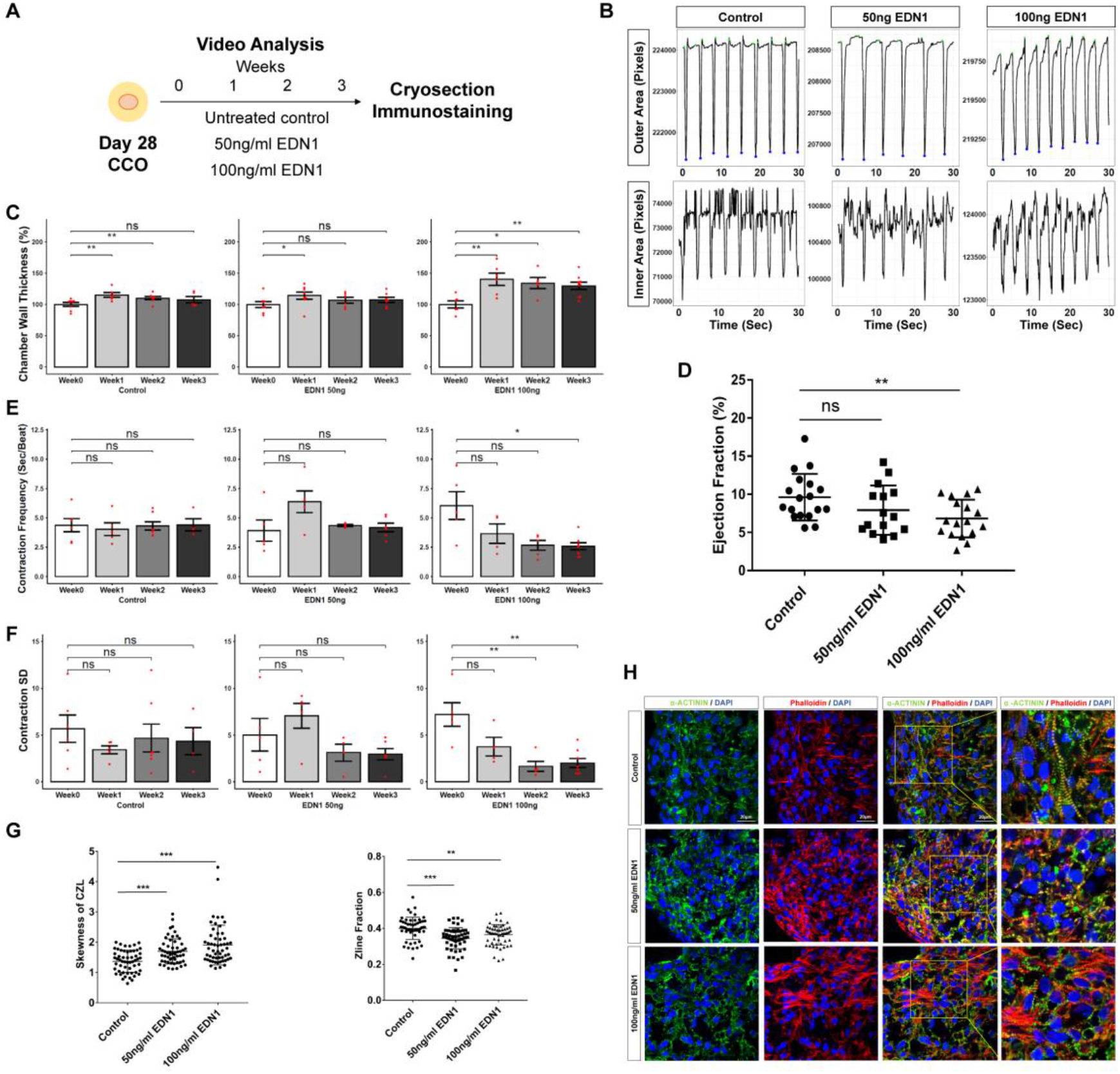
Cardiac hypertrophic phenotypes are recapitulated in chambered cardiac organoids treated with CH inducer EDN1. (**A**) Schematic diagram for cardiac hypertrophy modelling. (**B**) Outer (top) or inner (bottom) area against time plot of representative videos from a single CCO of each treatment condition. (**C, E and F**) Changes in individual analyzed parameters across the time points. Each chart represents one treatment condition (Left: Control; Middle: 50ng/ml EDN1; Right: 100ng/ml EDN1). Each red dot represents one organoid sample analyzed (n = 4-6 organoids per condition; mean ± s.e.m; Mann-Whitney U test). (**C**) Analysis of Chamber Wall Thickness normalized to Week 0. (**D**) Blind analysis of ejection fractions in CCOs on the third week of treatment showed significant reduced ejection fraction in 100ng/ml treated CCOs (Control n = 18, 50ng/ml EDN1 n = 15, 100ng/ml EDN1 n = 17; mean ± s.e.m; Welch’s t-test). (**E**) Analysis of Contraction Frequency. (**F**) Analysis of Contraction Standard Deviation. (**G**) Zline skewness (left) and Zline fraction (right) parameters are significantly affected in treated CCOs (n = 55 sections per group; mean ± s.e.m, Welch’s t-test). (**H**) Representative confocal immunofluorescence for z-line analysis of each condition at the end of Week 3. Scale bars, 20µm. (**C**-**H)** \*\*\**P* < 0.001; \*\**P* < 0.01; \**P* < 0.05.

To functionally characterize the CCO contractile machinery, the sarcomeric architecture of CMs within the CCO was assessed using the computational structural assay ZlineDetection (*26*). Earlier work demonstrated increased myofibrillar disarray in hPSC-CMs from CH patients (*24*). Therefore, CCO cryosections stained with phalloidin and α-actinin were computationally assessed to compute the fraction of α-actinin composed of well-formed z-lines and the skewness of the continuous z-line (CZL) (Fig. 4, G and H). Increasingly skewed CZL was observed as EDN1 treatment dosage increased, and significantly reduced fraction of Z-lines were well formed in both EDN1 treated groups, both highlighting the myofibrillar disarray induced by EDN1 treatment (Fig. 4G). These functional data strongly correlate with clinical observations of patients undergoing CH. CCOs were affected by the oxidative stress from the EDN1 treatment which have been shown to promote post-translational modification in sarcomere and myofilament proteins, altering the actin-myosin protein interactions and causing myofibrillar disarray (*27*). The disarray of the contractile apparatus sets the stage for myocardial energy depletion triggering adaptive responses. The CCOs exhibited these adaptive mechanisms by remodeling its contractile machinery, thickening the heart walls to counteract increased ventricular wall stresses, as well as increasing contraction rate to compensate for increased energy demands and reduced ejection fractions (*28*).

In conclusion, CCOs are able to maintain and mature the various cardiovascular subtypes in 3D and mimic the function of an *in vivo* heart and output important parameters such as heart wall thickness and ejection fraction. In this case, the heart wall thickness measured in CCOs is an important parameter for CH that is typically measurable only in an *in vivo* system. In addition, by tracking the contraction rhythm of individual CCOs, it is also possible to identify arrhythmic tendencies such as sinus tachy-/brady-cardias or more complex rhythmic disorders like atrial/ventricular fibrillations or torsades de pointes. Therefore, CCOs are demonstrated to be a useful *in vitro* platform that can complement current *in vivo* models to study adult cardiovascular diseases and conduct cardiotoxicity screening, with the ability to unveil potentially adverse mechanical and rhythmic alterations at high-throughput.

## Acknowledgments

The authors would like to thank Senior Consultant Ronald C.H. Lee from YLLSoM and NUHC Singapore for proofreading the manuscript and providing clinical perspective to the paper.

## Author contributions

BXH performed experiments; acquired, interpreted and analyzed the data; developed and validated the protocols described; generated the figures; and wrote and edited the manuscript. JKSP performed experiments; acquired, interpreted and analyzed the data; developed and validated the protocols described; generated the figures; performed bioinformatics analysis; and wrote and edited the manuscript. QHP, LCL, BMP contributed to cell culture and experiments. YC and YHL performed the organoid dissociation and scRNA-seq processing. OA and HHY contributed to the scRNA-seq bioinformatics analysis. VPS and JLYK designed the organoid video analysis pipeline. WKC contributed to experimental design and critical revision of the manuscript. SYN and BSS designed and conceived the study; interpreted the data; and wrote and edited the manuscript. All of the authors approved the manuscript.

## Data and materials availability

The data discussed in this publication have been deposited in NCBI’s Gene Expression Omnibus and are accessible through GEO Series accession number GSE168464 (https://www.ncbi.nlm.nih.gov/geo/query/acc.cgi?acc=GSE168464).

## Supplementary Materials

### Materials and Methods

#### Pluripotent stem cell lines and organoid culture

The human H7 hES cell line were cultured in feeder-free conditions on culture plates coated with Matrigel® matrix (Corning, U.S.A.); maintained in StemMACS™ iPS-Brew XF medium (Miltenyi Biotec, Germany). Culture medium was changed daily. After 80–90% confluency was reached, hES were passaged using ReLeSR (STEMCELL Technologies). At 90-95% confluency, differentiation was started using RPMI-1640 Medium (1x) (Hyclone) with B-27 Minus Insulin 50x supplement (Gibco, U.S.A.) and 1% Pen/Strep (Gibco, U.S.A) containing 12µM CHIR99021, a GSK inhibitor (Miltenyi Biotec), as described by Lian et al., 2013 (*17*). After 24 hours in culture, the medium was replaced with RPMI/B27 minus insulin for 2 days. On day 3, the cells were cultured in the same basal media containing an inhibitor of WNT signaling, such as IWP-2 (Miltenyi Biotec). On day 5, the medium was replaced with the same basal media. On Day 7, the cells were dissociated into single cells using StemPro-Accutase®(Gibco, U.S.A.). The cells were resuspended in CGM media and 500k cells were seeded into each well of a 96 well, ultra-low attachment round bottom plate (CoSTAR, 2022-03-14). The plate was subsequently spun down at 1000rpm for 3min and the supernatant was removed from each well. With 2x Matrigel® matrix (Corning, U.S.A.) diluted in DMEM/F12 (Gibco, U.S.A.), 20 µl of matrigel was added into each well, followed by a quick spin at 1000rpm for 2min. The 96 well plate was incubated at 37 °C under humidified atmosphere of 5% CO2 for an hour. After one hour, 100 µl of CGM media (B27+ supplement, 17.5ng/ml Thrombin, 3ng/ml Cardiotrophin-1, 2ng/ml EGF, 5ng/ml FGF-2, 20ng/ml WNT-3A, 20ng/ml EDN1) was added into each well, drop-by-drop. Fresh media was changed every two days. On day 14, organoids were transferred from the 96 well plates into a spinner flask containing 50ml of RPMI/B27 media supplemented with 1% Pen/Strep, and placed on a spinner platform at 60 rpm.

#### Fluorescence-activated cell sorting and flow cytometric analysis

On day 7 of the cardiomyocyte differentiation, the cells were dissociated using StemPro-Accutase®(Gibco, U.S.A.) and stained with EDNRA antibody (1:200 dilution, Invitrogen; #MA3-005) for 2 h at 37 °C in blocking buffer (1% FBS in PBS). After 2 h, the cells were washed and stained with anti-human CD172a/b (SIRPα/β) antibody conjugated antibody (1:250 dilution, Biolegend; #323808) and donkey anti-rabbit Alexa Fluor® 594 secondary antibody (1:1000 dilution, Invitrogen; #R37119) for 1 hour. The cells were washed thoroughly and resuspended in FACS buffer (0.5% FBS, 1% BSA in PBS) and FACS was performed using LSR II (BD Biosciences, U.S.A.).

#### Immunohistochemistry

Organoids were washed in PBS before fixing in 4% PFA and placed on a rotator in 4°C overnight. The excess fixative was removed with three washes of PBS and organoids were then incubated in a 15% sucrose solution in 4°C overnight, followed by 30% sucrose in 4°C overnight. Finally, the organoids were embedded in OCT for snap-freezing on dry ice. For immunohistochemical stainings, organoids were sectioned in slices of 10µm thickness at -15 to -20°C. Then, sections were stored in -70°C for long term storage. For immunostaining, organoid sections were washed with PBS thrice and the tissue was fixed with 4% PFA for 10min, followed by permeabilizing solution (0.1% Triton-X in PBS) for 10min. Afterwards, sections were incubated overnight in 4°C in blocking buffer (5% FBS in PBS) containing antibodies anti-cTnT (rabbit,1:400, Abcam, ab45932), anti-ISL1 (rabbit, 1:250, Abcam, ab109517), anti-CleavedCaspase-3 (rabbit, 1:400, CST, D175) anti-KI67 (mouse, 1:800, CST, 8D5), anti-Alpha-actinin (mouse, 1:100, Abcam, Ab9465). On the next day, sections were washed thrice with PBS and incubated for 1-1.5 h at room temperature in secondary antibody in Alexa Fluor® 594-conjugated anti-rabbit antibody (1:1000), Alexa Fluor® 594-conjugated anti-mouse antibody (1:1000), Alexa Fluor® 488-conjugated anti-rabbit antibody (1:1000), Alexa Fluor® 594-conjugated anti-mouse antibody (1:1000). Stained organoid cryosections were imaged using a confocal laser scanning Olympus Fluoview FV1000 microscope using 10x, 20x and 100x magnification objectives. The images were then stacked and further processed using Olympus FV10-ASW 4.2 Viewer.

#### Magnetic bead sorting of cardiomyocyte

hES-derived cardiomyocytes and cardiac organoids were dissociated with accutase, cells were blocked with a solution containing 0.5% bovine serum albumin (BSA) and 2mM EDTA. Using human PSC-derived cardiomyocyte isolation kit (Miltenyi Biotec, Germany), cardiomyocytes were isolated using a two-step isolation protocol as per the manufacturer’s instructions.

#### RNA Extraction

For cultured cell samples, cells were collected and lysed in 300 µl of TRIzol reagent (Invitrogen, U.S.A.). The samples were allowed to stand for 5 min at room temperature, after which 180 µl of Chloroform (Kanto Chemical, Japan) was added to allow for phase separation by centrifugation at 12,000 × *g* for 15 min at 4 °C. Next, aqueous phase was transferred to a fresh tube with equal volumes of isopropanol and GlycoBlue Coprecipitant (Invitrogen, U.S.A.). The samples were incubated at room temperature for 20 min. The samples were pelleted through centrifugation at 12,000 × *g* for 15 min at 4 °C. The RNA pellet was washed with 100% ethanol, air-dried before reconstituting it in nuclease-free water (Ambion, U.S.A.).

#### Reverse transcription and quantitative real-time PCR

RNA samples (250-500 ng) were reverse transcribed to obtain cDNA using High-Capacity cDNA Reverse Transcription Kit (Applied Biosystems, U.S.A.). qPCR was performed using the FAST SYBR Green Mix (Applied Biosystems, U.S.A.), 0.3 μM of specific primers (Table S1) and ∼5 ng of cDNA. ΔΔCT-based relative quantification method was adopted for qPCR analysis using the QuantStudio 5 384-well Block Real-Time PCR system (Applied Biosystems, U.S.A.). The threshold cycle was determined to be ≥35. Data is presented as fold-change where CT values were normalised to *β-ACTIN*. Data presented are representative of three independent experiments with error bars indicative of the standard deviation unless otherwise stated.

#### Organoid Dissociation

Organoids of various ages (Day 8, Day 14, and Day 21) were dissociated using TrypLE (ThermoFisher, 12604013). Organoids were washed once with 1xPBS and then incubated in TrypLE solution at 37 °C for 10 to 15 mins. Subsequently, organoids were dissociated into single cells by gentle pipetting using wide-bore tips. Cell solutions were then diluted with 1xPBS and passed through 40um strainer (Falcon, 352340) to remove any undissociated cell clumps. Cells were then centrifuged down to remove TrypLE and washed once with 1xPBS (300 × *g* for 5 min at 4 °C). Pellets were eventually re-suspended in 1xPBS and stored on ice.

#### 10x Genomics single-cell RNA Library preparation and sequencing

Cell viability and concentration were checked using an automated cell counter (BioRad). Cell stocks were diluted to around 1000 cells/uL with 1xPBS before downstream steps.

Chromium™ Single Cell 3’ Library & Gel Bead Kit v2 was used to construct the library. For all library preparation, recovery of 6000 cells was aimed. Briefly, cells together with Gel Bead-in-Emulsions (GEM)-reverse transcription reagents, and GEM beads were loaded to the 10xChip, in which individual cells are captured, lysed, reverse transcribed and barcoded.

Subsequently, barcoded cDNA was amplified and used for library construction. Libraries which passed the quality control were then sent out for sequencing.

#### Bioinformatics

Cell filtering, data normalization and unsupervised analysis were carried out using R library *Seurat* version 3.2.1 (*29, 30*). The cells were filtered based on their number of gene features and percentage of mitochondrial genes. The threshold used for gene features is between 2000 to 7500, and less than 10% mitochondrial gene expression. Genes to be used in analysis were expressed in at least 10 cells. LogNormalize function was used to normalize gene expression within each cell, by dividing the counts of each gene by the total counts in the cell, followed by a scale factor multiplication of 10,000 and then a natural log-transform of the result.

Variable genes were identified using the FindVariableFeatures function and used to perform a principal component (PC) analysis using RunPCA. Within all the PCs, we used the top 20 PCs to do Uniform Manifold Approximation and Projection (UMAP) analysis. Clustering was carried out using the FindClusters function. Gene expression differences between the various cell clusters was carried out using FindAllMarkers function, filtering out genes expressed in less than 25% of cells in the compared clusters, and selecting genes with differential expression levels above 25%. Finally, clusters were identified by going through the list of differentially expressed genes and comparing the highly expressed genes with published literature.

#### Video analysis

Video clips of the organoids were imaged using Nikon ECLIPSE Ti-S fluorescent microscope and recorded using an Andor Zyla 4.2 sCMOS. Images were extracted from each video clip at a frequency of 1/10^th^ second using ffmpeg (*31*), generating approximately 300 images from each video. The images were analysed using an image analysis pipeline implemented using Cell Profile 3.1.9 (*32*). Organoid boundary and inner chamber regions were independently segmented from the images and used to compute the organoid size (outer area), the inner chamber area, the heart wall area and the heart wall area as a fraction of the organoid area (fig. S1). The organoid contraction peaks were computed in R by taking the first order differential maximas of the organoid area with respect to time. The contraction peaks were then used to compute the individual organoid contraction rate and variations by taking the mean and standard deviation of the contraction intervals respectively.

#### Statistical analysis

Statistical analyses were performed either using GraphPad Prism 8 or R. Quantitative PCR values between groups are compared using two-tailed Student’s t-tests. Cardiac Organoid parameters comparison between groups were done using two-tailed Mann Whitney U-test when sample size is small, otherwise Welch’s t-test is used.

**Fig. S1.**
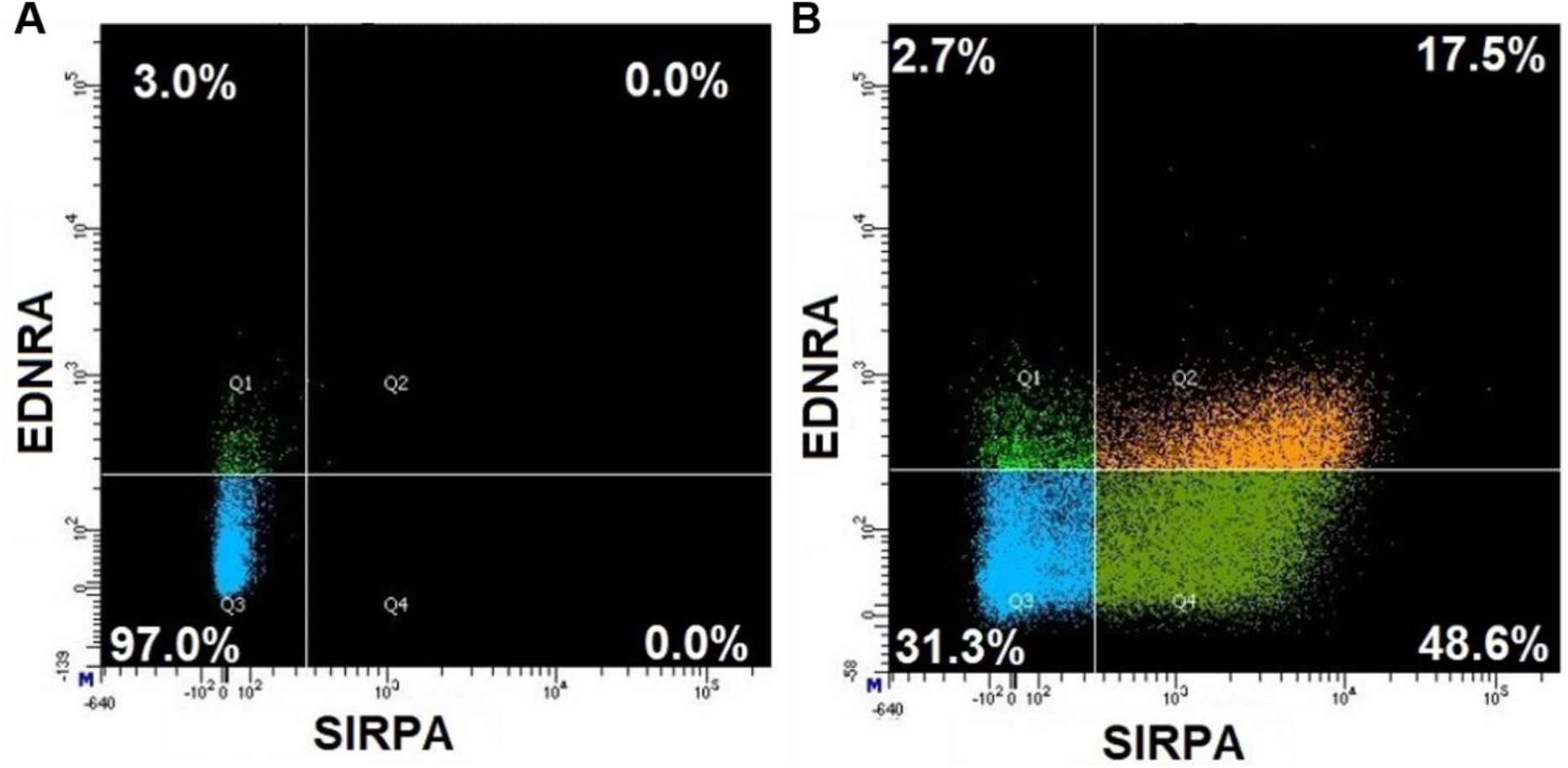
Sorting parameters used to obtain individual cell populations to generated chambered cardiac organoids of varying compositions. **(A)** Unstained, mixed control population of day 7 differentiated iPSC-cardiomyocytes. **(B)** Same mixed, control population stained with SIRPα and EDNRA fluorescent antibodies. SIRPα single-positive cells (bottom right quadrant) contain the Cardiomyocyte population while the SIRPα, EDNRA double-positive cells (top right quadrant) contain the Cardiovascular Progenitor population.

**Fig. S2.**
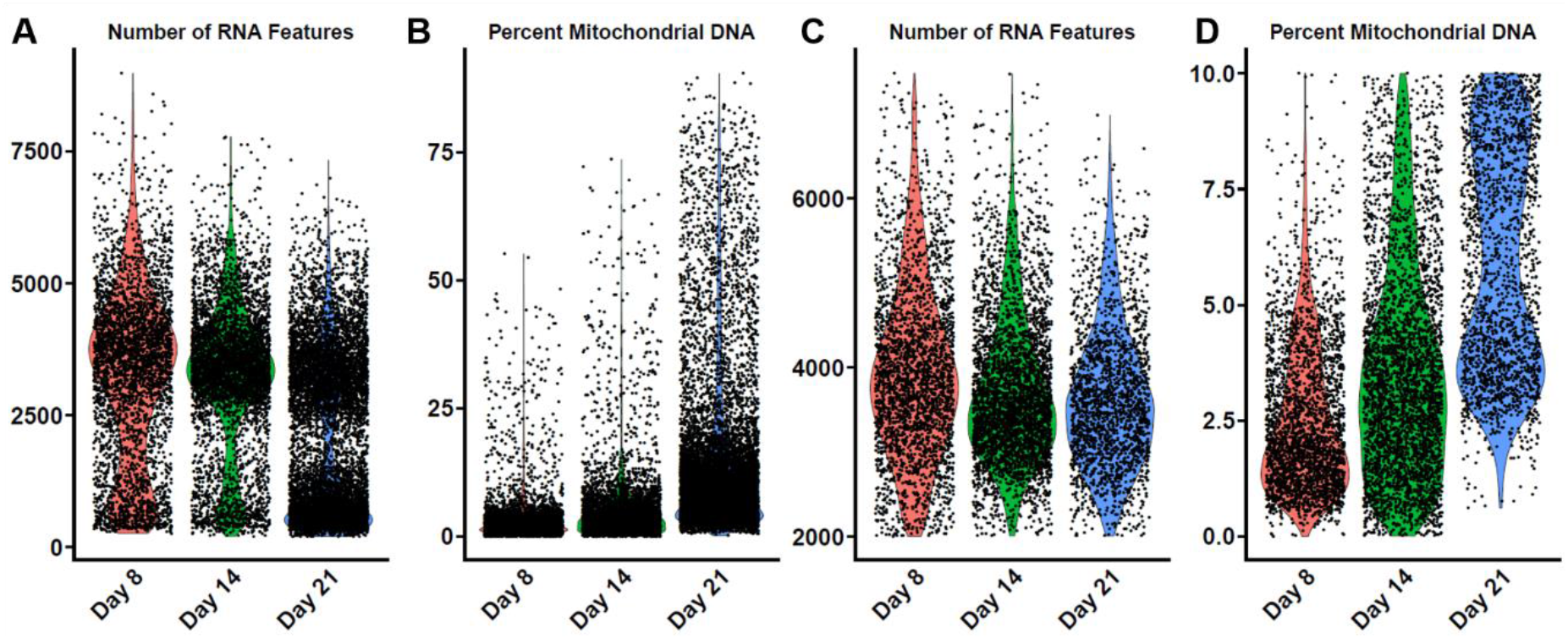
Quality control filtering of the dataset to only include true cells of high quality using two metrics – Number of RNA Features and levels of mitochondrial gene expression. **(A)** Unfiltered RNA gene detection levels across all single cell reads. **(B)** Unfiltered percentage of mitochondrial DNA detected within all single cell reads. **(C)**, Filtered RNA gene detection levels across all single cell reads, keeping cells that express between 2000 – 7500 unique genes. **(D)**, Filtered percentage of mitochondrial DNA detected within all single cell reads, keeping cells that have <10% mitochondrial DNA.

**Fig. S3.**
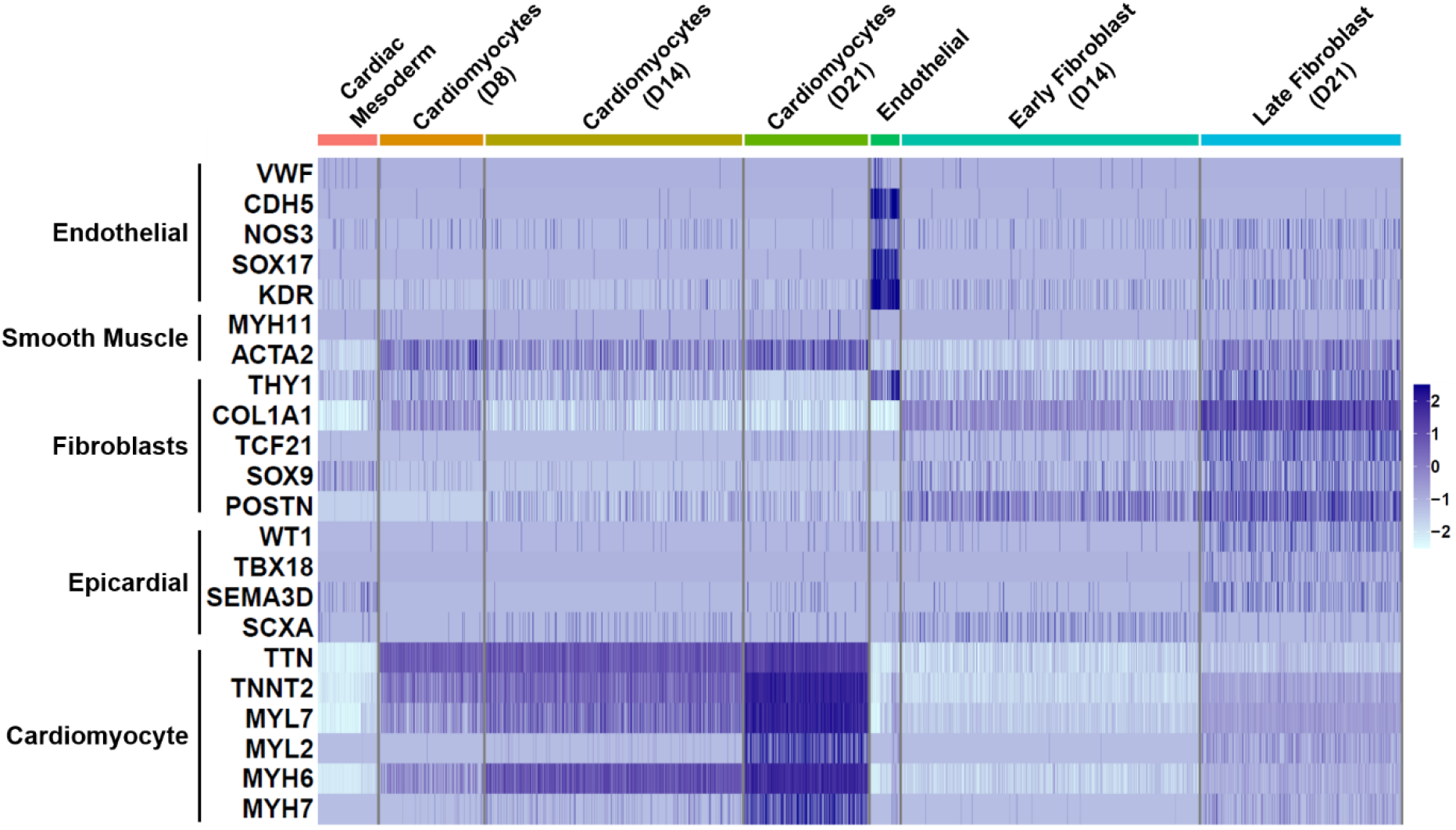
Heatmap depicting the varying expression levels of marker genes of each identified cluster. Gene expression levels of the seven clusters related to the cardiac lineage are isolated in this heatmap. Each cluster identified are expressing their own marker genes.

**Fig. S4.**
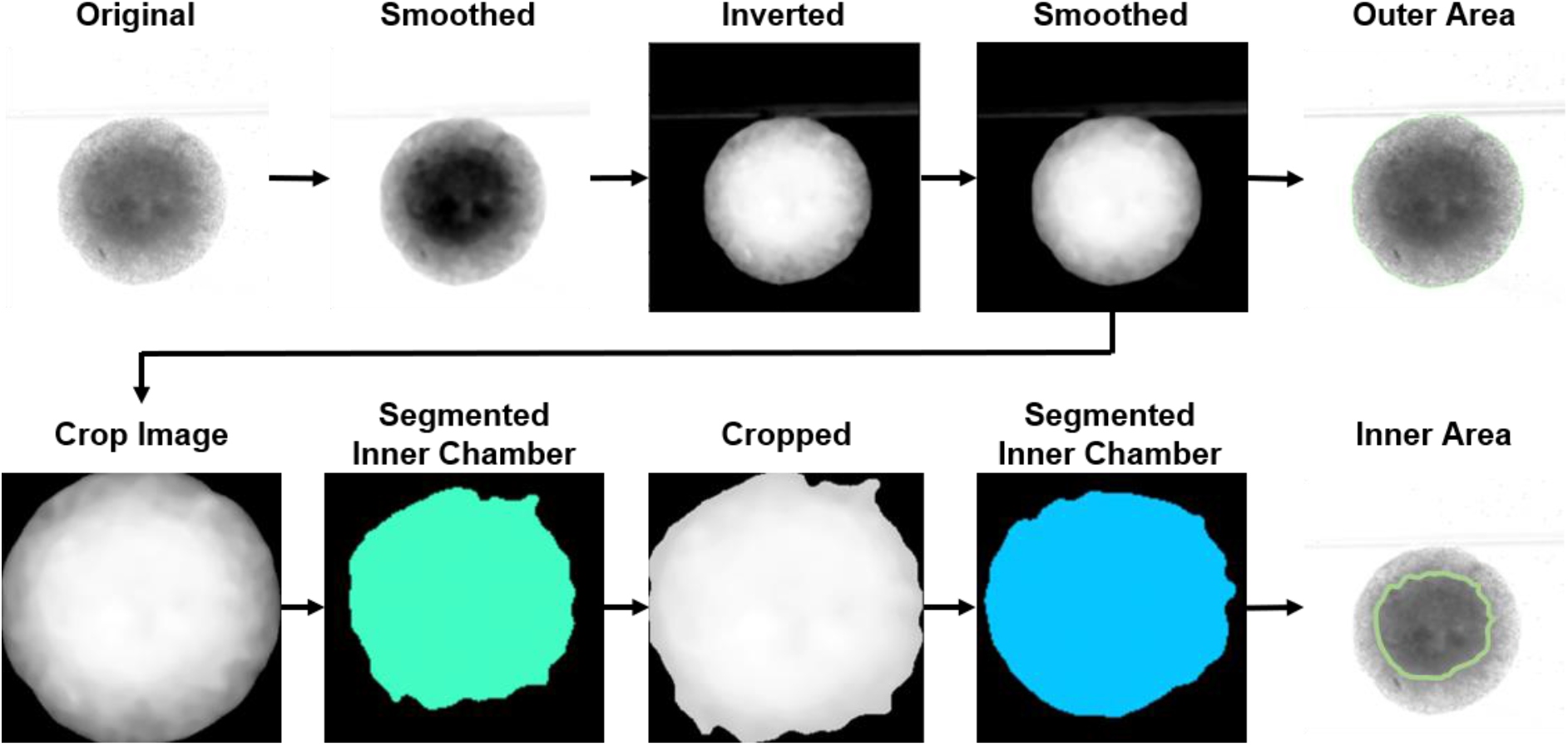
Schematic depicting the pipeline of image segmentation. Individual frames of the organoid videos were smoothened to allow segmentation of the organoid and inner chambers.

**Table S1.**
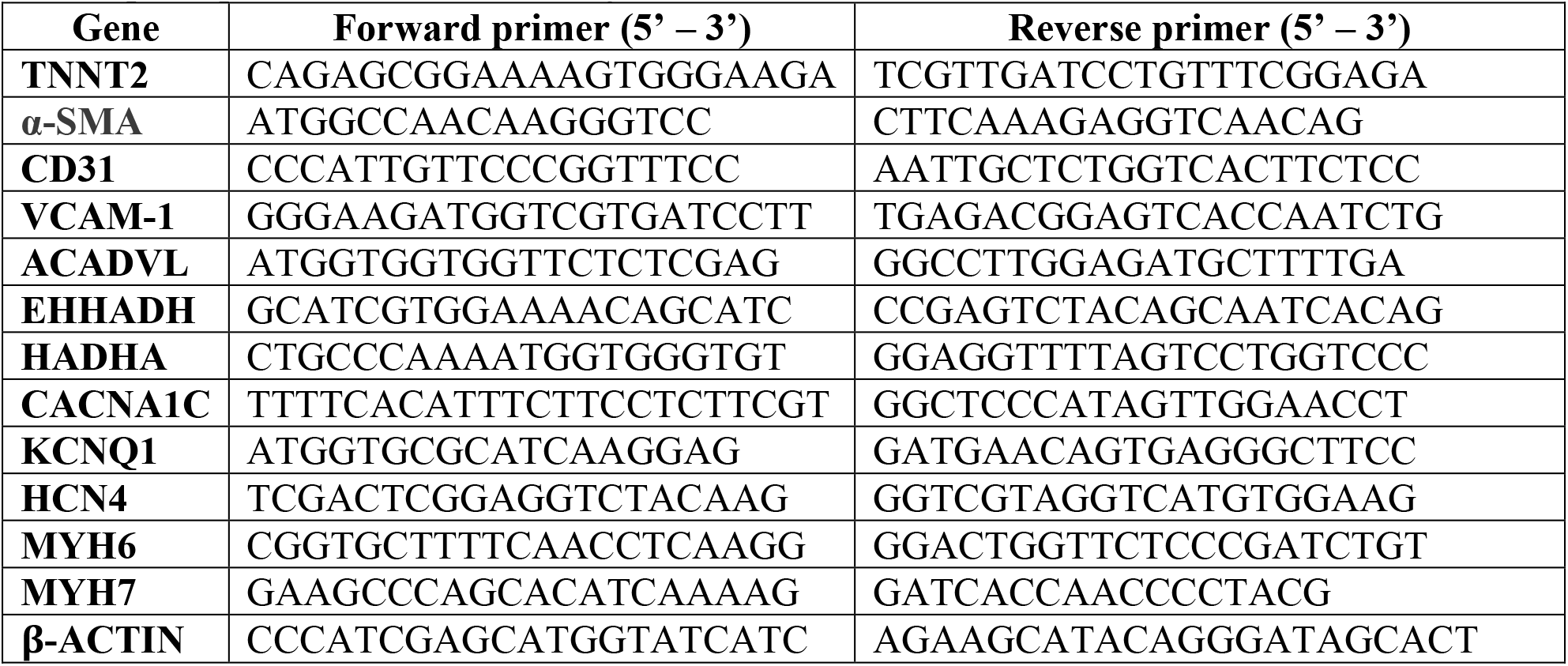
List of qPCR primers used in this study

**Movie S1.**

Representative brightfield video tracking the contractions of Week 3 untreated control Chambered Cardiac Organoid.

**Movie S2.**

Representative brightfield video tracking the contractions of Week 3 post 50ng/ml EDN1 treatment Chambered Cardiac Organoid.

**Movie S3.**

Representative brightfield video tracking the contractions of Week 3 post 100ng/ml EDN1 treatment Chambered Cardiac Organoid.

**Data S1.**

Table of top 100 differentially expressed genes upregulated between clusters identified from the single cell RNA sequencing data across the three timepoints of organoids. Annotations retrieved from Ensembldb database.

